# Aurora kinase A is essential for meiosis in mouse oocytes

**DOI:** 10.1101/2021.01.08.425851

**Authors:** Cecilia S. Blengini, Patricia Ibrahimian, Michaela Vaskovicova, David Drutovic, Petr Solc, Karen Schindler

**Affiliations:** Department of Genetics; Rutgers, The State University of New Jersey; Human Genetics Institute of New Jersey; Institute of Animal Physiology and Genetics of the Czech Academy of Sciences, Libechov, Czech Republic

**Keywords:** Aurora kinase, AURKA, meiosis, oocyte, microtubule organizing center, fertility

## Abstract

The Aurora protein kinases are well-established regulators of spindle building and chromosome segregation in mitotic and meiotic cells. In mouse oocytes, there is significant Aurora kinase A (AURKA) compensatory abilities when the other Aurora kinase homologs are deleted. Whether the other homologs, AURKB or AURKC can compensate for loss of AURKA is not known. Using a conditional mouse oocyte knockout model, we demonstrate that this compensation is not reciprocal because female oocyte-specific knockout mice are sterile and their oocytes fail to complete meiosis I. In determining the AURKA-specific functions, we demonstrate that its first meiotic requirement is to activate Polo-like kinase 1 at microtubule organizing centers (MTOCs; meiotic spindle poles). This activation induces fragmentation of the MTOCs, a step essential for building a bipolar spindle. The next step that requires AURKA is building the liquid-like spindle domain that involves TACC3. Finally, we find that AURKA is also required for anaphase I onset to trigger cohesin cleavage in an APC/C independent manner. We conclude that AURKA has multiple functions essential to completing MI that are distinct from AURKB and AURKC.

**Author Summary:** Female gametes, oocytes, are uniquely prone to chromosome segregation errors in meiosis I that are associated with early miscarriages. The Aurora protein kinases are essential to control chromosome segregation in all cell types. During mitosis, Aurora kinase A (AURKA) regulates the building of the spindle, the machinery responsible for pulling chromosomes apart. Here, we use a genetic approach to demonstrate that AURKA is essential for meiosis I in mouse oocytes. AURKA is required at multiple steps in meiosis I, first to trigger fragmentation of protein structures that make up the two ends of the meiotic spindle, later to regulate building of a specialized phase-separated spindle domain, and finally to trigger efficient cleavage of cohesin, the molecular glue that holds chromosomes together until anaphase onset. These findings are the first demonstration of distinct Aurora kinase function that cannot be compensated for by the other two homologs. Therefore, this mouse model is excellent tool for pinpointing specific Aurora kinase functions and identifying AURKA target proteins critical for chromosome segregation in meiosis I.

## Introduction

Haploid gametes, which are required for sexual reproduction, are generated through meiosis; a cell division that undergoes two successive rounds of chromosome segregation without an intervening round of DNA replication. First, homologous chromosomes are separated during meiosis I (MI), followed by sister chromatid separation in meiosis II (MII). Errors in MI give rise to aneuploid gametes that, if fertilized, lead to congenital birth defects or embryo development failure [1–3]. Critical to accurate chromosome segregation is the formation of a bipolar spindle apparatus which captures chromosomes and pulls them apart. Therefore, defects in spindle building could cause chromosome mis-segregation and aneuploidy.

In somatic cells, spindles are built from microtubules that nucleate from centrosomes. Centrosomes are cellular structures that form the ends, or poles, of the spindle and are composed of centrioles surrounded by organized layers of pericentriolar material (PCM). However, in mammalian oocytes this process is strikingly different because centrioles were lost during oocyte development [4]. In mouse oocytes, spindle formation depends on multiple microtubule organizing centers (MTOC) that lack centrioles but retain PCM that nucleate microtubules [5–9]. During spindle formation, MTOCs undergo a series of highly regulated, morphological changes. First, MTOCs decondense and fragment into smaller MTOCs. Next, these small MTOCs are sorted so that after an intermediate multi-polar ball-like formation, they finally cluster into the two poles of the spindle [9, 10]. Perturbation of any of these steps dramatically affects the spindle structure and the interaction between microtubules and chromosomes, which ultimately can alter chromosome segregation. One result of this perturbation is that oocytes fail to complete meiosis because they activate the spindle assembly checkpoint (SAC) that monitors attachment of microtubules to kinetochores and delays anaphase onset until kinetochores are appropriately attached [11].

The Aurora kinases (AURK) are a family of serine/threonine protein kinases involved in chromosome segregation, in mitosis and in meiosis [12–14]. This protein family has three members: AURKA, AURKB and AURKC. Most somatic cells express only AURKA and AURKB, but oocytes express all three isoforms. In somatic cells, AURKA localizes to centrosomes and is involved in centrosome maturation and separation [15–17] and microtubule nucleation [18, 19]. However, in meiosis, two AURKs are needed to build a normal bipolar spindle: AURKA and AURKC [20]. AURKA localizes to MTOCs [21–23], and may contribute to spindle formation through mechanisms different than those used in mitosis: regulating MTOC numbers [21, 22, 24], the distribution of MTOCs into two poles [10, 22] and maintaining spindle pole structure [25, 26]. Furthermore, AURKA activity is required to assemble a liquid-like spindle domain (LISD) composed of several regulatory factors. The LISD is proposed to allow rapid, and localized, protein concentration changes around microtubules during spindle formation [8]. Depletion or inhibition of AURKA in mouse oocytes produces short, disorganized spindles, characterized by over-clustered MTOCs and loss of the LISD [8, 21, 23, 26]. Consistent with these spindle abnormalities, these depleted or inhibited oocytes fail to complete meiosis and arrest in metaphase I [21, 22]. AURKC also localizes to MTOCs and contributes to MTOC clustering into two spindle poles. Prevention of AURKC from localizing to MTOCs in mouse oocytes causes frequent multipolar spindle formation and increased rates of aneuploid egg production [20].

In oocytes, the AURKs exhibit complex genetic interactions and compensatory abilities. For example, AURKB is the catalytic component of the chromosome passenger complex (CPC) in mitosis. But, in oocytes, AURKC outcompetes AURKB and takes over this CPC role [27, 28]. Furthermore, oocytes can complete meiosis in the absence of both AURKB and AURKC because AURKA can function in the CPC; this is specific to oocytes because this compensation does not occur in HeLa cells or in spermatocytes [29, 30]. However, although AURKA can compensate, it is not complete because a subset of oocytes arrest in metaphase I with short spindles. These short spindles arise because AURKC is required outcompete AURKA from CPC-binding to keep AURKA at MTOCs and ensure appropriate spindle length [29]. Because AURKA and AURKC compete for CPC binding and because a second population of AURKC exists at MTOCs, we asked if the compensatory abilities of AURKA and AURKC were reciprocal.

To test if AURKC can compensate for loss of AURKA, and to further understand the role of AURKA during meiosis in mouse oocytes, we generated a mouse strain that lacks *Aurka* [16] specifically in oocytes using *Gdf9*-mediated Cre excision [31]. Consistent with AURKA being the most abundant AURK in oocytes [29], we demonstrate that AURKA is essential for oocyte maturation through fragmenting MTOCs, building the LISD and triggering cleavage of cohesin in an APC/C-independent manner. Moreover, we demonstrate that that AURKB and AURKC cannot compensate for loss of AURKA. Therefore, AURKA is the only Aurora kinase essential for MI in mouse oocytes.

## RESULTS

### Generation and confirmation of mice lacking *Aurka* in oocytes

Because AURKA can function in the CPC in the absence of AURKB and AURKC [29], we asked if similar compensatory functions exist in the absence of AURKA. Prior AURKA studies used small-molecule inhibitors such as MLN8237 and overexpression to investigate AURKA’s role in mouse oocyte meiotic maturation which do not allow for compensation studies [8, 21, 22, 24, 26, 29, 32]. To assess compensation and potential AURKA-specific requirements, we deleted *Aurka* (*Aurka*^*fl/fl*^) using *Gdf9-*Cre. *Gdf9* expression begins around day 3 after birth in prophase I-arrested oocytes; these oocytes already completed early prophase I events such as chromosome synapsis and recombination. *Aurka* is therefore deleted in growing oocytes prior to completion of chromosome segregation in meiosis I. To confirm that AURKA was depleted from oocytes, we first assessed total AURKA levels by Western blotting. When normalized to the AURKA signal in oocytes from wild-type (WT; *Aurka*^*fl/fl*^) littermates, the signal in *Aurka* knockout (KO; *Aurka*^*fl/fl*^ *Gdf9-Cre*) oocytes was at background levels (Fig. 1A-B). We also assessed the presence of AURKA at Metaphase I (Met I) by immunocytochemistry. In WT oocytes, AURKA localized to Met I spindle poles. Compared to WT, *Aurka* KO oocytes lacked AURKA signal (Fig. 1C-D). Finally, we measured the activity of AURKA by immunostaining oocytes with anti-phosphorylated CDC25B-serine 351 (pCDC25B), an AURKA substrate that localizes to spindle poles [33]. Consistent with the loss of polar AURKA, there was no detectable pCDC25B in *Aurka* KO oocytes (Fig. 1E-F). These data indicate that Gdf9-mediated Cre excision of *Aurka* is sufficient to deplete AURKA in mouse oocytes.

**Fig. 1.**
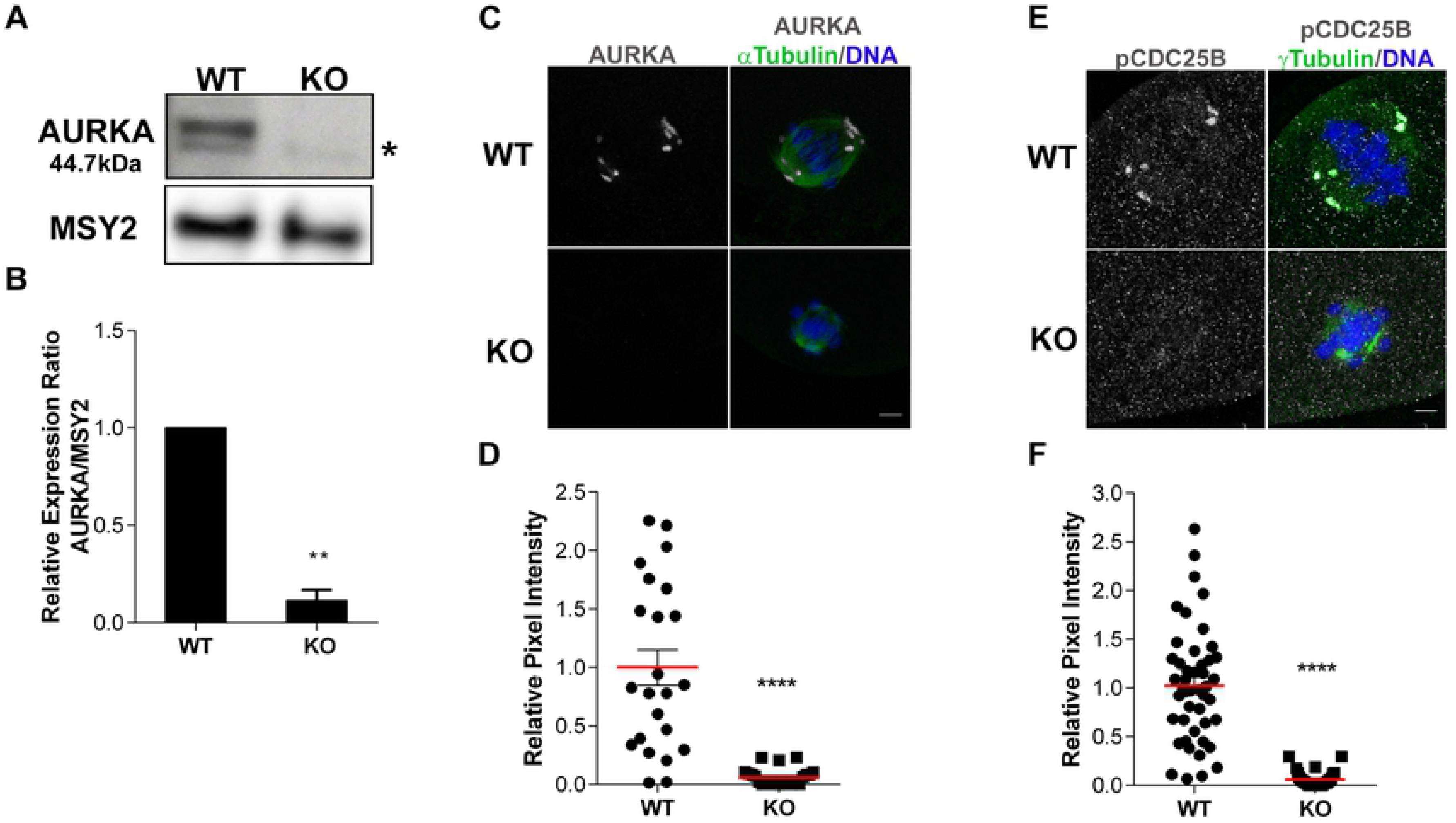
AURKA is deleted from oocytes. **(A)** Western blot detecting AURKA from prophase-I arrested wild-type (WT) and *Aurka* knockout (KO) oocytes (100 oocytes/lane). After stripping the membrane, MSY2 served as loading control. Bands at ~43kDa were included in the quantifications for AURKA signal. n=4 animals/genotype/experiment. Asterisk = non-specific band **(B)** Quantification of AURKA after normalizing to MSY2 in (A) (Unpaired Student’s t-Test, two-tailed, ** p<0.0035) **(C-F)** Localization and activity of AURKA in WT and KO oocytes. **(C)** Representative confocal images of metaphase I oocytes immunostained with antibodies against AURKA (gray), α-Tubulin (green) and DAPI (blue); **(D)** Quantification of AURKA intensity in (C); Unpaired Student’s t-Test, two-tailed, **** p<0.0001; number of oocytes, WT: 23; KO: 24). **(E)** Representative confocal images of metaphase I oocytes immunostained with antibodies against phosphorylated CDC25B (gray; pCDC25B), γ-Tubulin (green) and DAPI (blue). **(F)** Quantification of pCDC25B intensity in (E); (Unpaired Students t-Test, two-tailed, **** p<0.0001; number of oocytes, WT: 46; KO: 30). Graphs show individual values plus the mean ± SEM from 2-3 independent experiments. Scale bars: 10μm.

### *Aurka*-oocyte knockout mice are sterile

To determine the consequence of deleting *Aurka* in mouse oocytes, we conducted fertility trials. Age-matched WT and KO females were mated to WT B6D2F1/J males of proven fertility and the numbers of pups born were recorded. We carried out this fertility trial for the time it took WT females to produce 5 litters (~4 months). Compared to WT females that produced ~6 pups/litter, *Aurka* KO females never produced a pup (Table 1). Therefore, in contrast to AURKB and AURKC [28, 29], oocyte expressed AURKA is essential for female fertility.

**Table 1.**
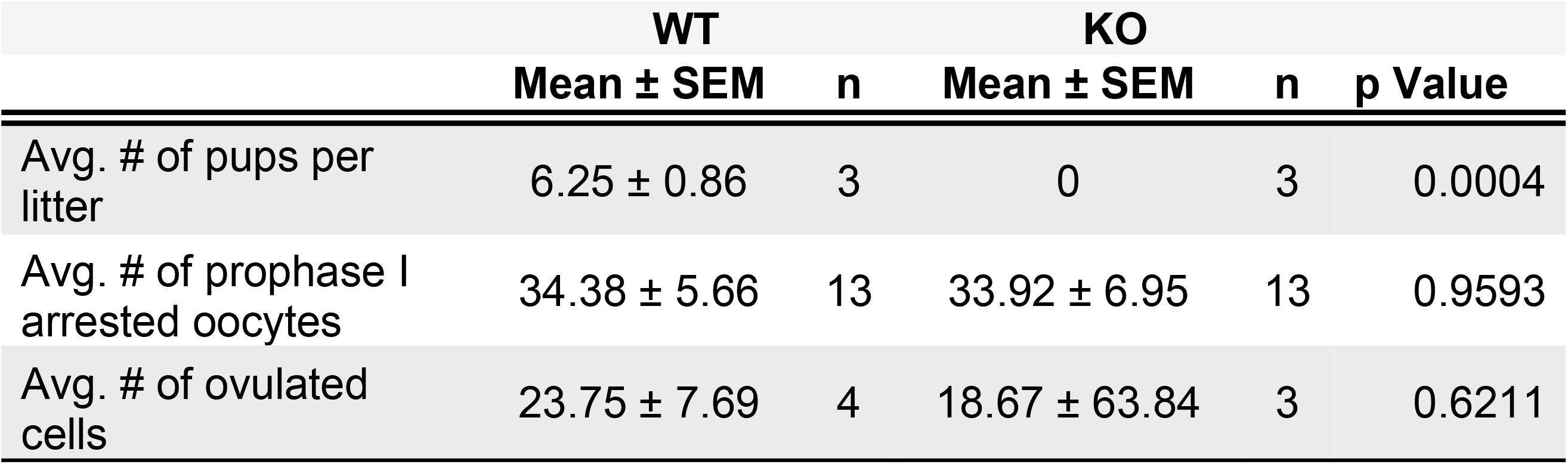
Number of pups, oocytes and cells ovulated from WT and *Aurka* KO females

To understand the cause of sterility, we first evaluated follicle development in histological sections of ovaries from females at different ages (1m, 2m, 6m). Of note, the animals used for histological sampling at 6 months were the females used the fertility trial. Compared to age-matched WT animals, there were no significant differences in the number of follicles at different developmental stages (Fig. 2A-F). Importantly, *Aurka* KO ovaries contained corpus luteum (CL), indicating that these females can ovulate. However, *Aurka* KO ovaries had 50% reduction in the number of CL in comparison to WT (Fig. 2C-F) suggesting that not all oocytes in fully developed follicles were ovulated. Taken together, these data indicate that the remaining Aurora kinases, AURKB and AURKC cannot compensate for loss of AURKA.

**Fig 2.**
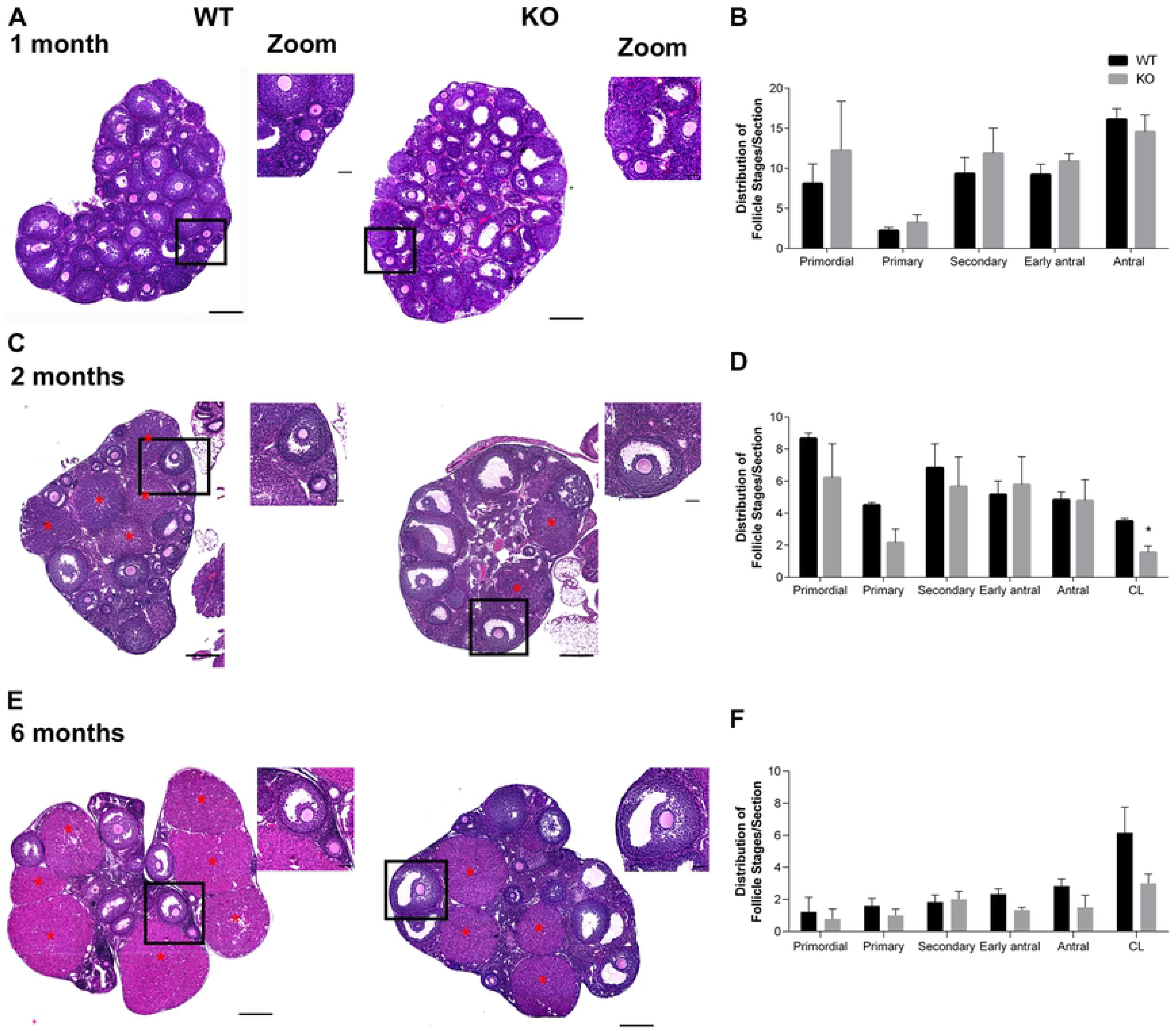
*Aurka* KO females have normal follicle development. **(A, C, E)** Representative images of hematoxylin/eosin-stained ovarian sections from WT and *Aurka* knockout (KO) females from different ages: 1 month **(A)**; 2 months **(C)**; 6 months **(E**), red asterisk: corpus luteum (CL). The zoom panels highlight commonly observed follicles at each age. **(B, D, F)** Quantification of follicle types from the ovaries represented in (A, C, E) respectively. Follicle numbers were quantified for each ovary and reported as the average number of each type of follicles per section. * p < 0.05. Graph shows the mean ± SEM (1 and 6 months: 3 females/genotype, 2 months: 2 WT; 3 A KO). Scale bars: 50μm (zoom panels) and 200μm.

### AURKA has unique functions during meiosis I

Because *Aurka* KO females are sterile but ovulate, we next assessed the quality of the ovulated cells. We induced ovulation through hormonal stimulation and harvested cells from oviducts. In this strain background, ~80% of cells in oviducts from WT mice contained polar bodies (Fig. 3A-B), indicating a completion of meiosis I (MI) and arrest at Metaphase of meiosis II (Met II). In contrast, none of the cells from KO oviducts had polar bodies, and they all were arrested in Met I, indicating a failure to complete MI (Fig. 3A-B). We note that similar number of cells were obtained from WT and KO oviducts (Table 1).

**Fig 3.**
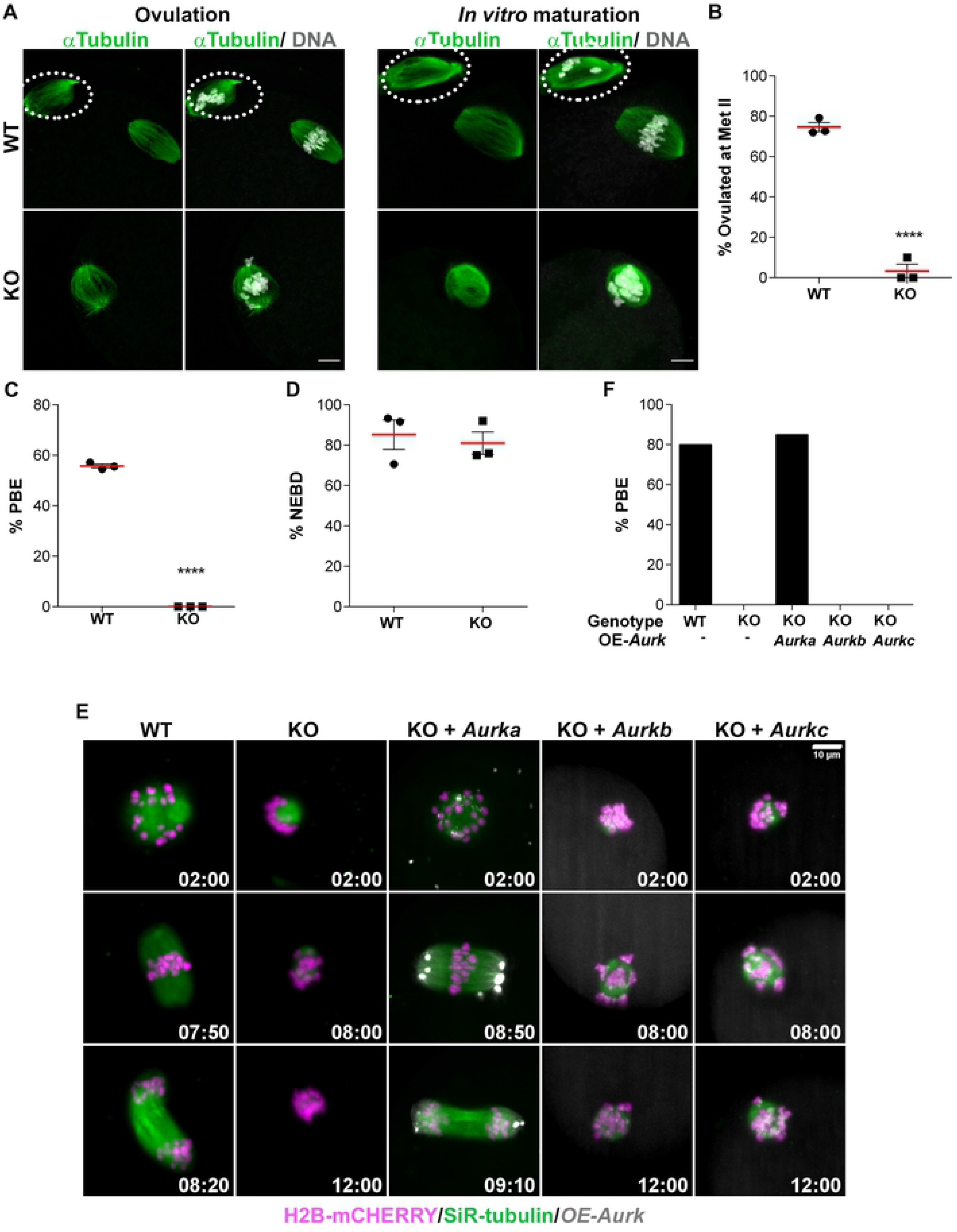
AURKA is specifically required in oocytes to complete meiosis I. **(A)** Representative confocal images of the oocytes and eggs retrieved from oviducts of WT and *Aurka* KO females or oocytes matured *in vitro.* Cells were immunostained with antibodies against α-Tubulin (green) and DAPI (gray); **(B)** Quantification of percentage (%) of cells ovulated at Metaphase II (Met II); (Unpaired Students t-Test, two-tailed, **** p<0.0001). **(C)** Quantification of the % of oocytes that undergo polar body extrusion (PBE) *in vitro* (Unpaired Students t-Test, two-tailed, **** p<0.0001). **(D)** Quantification of the % of oocytes that undergo nuclear envelope breakdown (NEBD) *in vitro* (Unpaired Students t-Test, two-tailed, p=0.6707). Graphs show the mean ± SEM from 3 independent experiments (3 females/genotype). Scale bars: 10μm. **(E)** Live light-sheet imaging of WT and KO oocytes expressing histone H2B-mCHERRY (magenta) and stained with SiR-tubulin (green). Some KO oocytes also expressed exogenous AURKA-EGFP, AURKB-EGFP or AURKC-EYFP (gray). Maximum intensity z-projections and selected time points are shown. Scale bars: 10 μm. (F) Data in (E) was used to quantify % of oocytes extruding polar body. n (WT, KO, KO + *Aurka*, KO + *Aurkb*, KO + *Aurkc*) = 10, 10, 13, 15, 11, respectively.

To identify where in MI *Aurka* KO oocytes were failing, we examined oocytes that were matured *in vitro* for a time in which WT oocytes would reach the Met II arrest. We isolated similar numbers of prophase I-arrested oocytes from WT and KO females, consistent with the ovarian reserve not being affected (Table 1). After maturation, WT oocytes extruded polar bodies *in vitro*. In contrast, none of *Aurka* KO oocytes extruded polar bodies (Fig. 3A, C). We did not observe a difference between WT and KO oocytes in the percentage of oocytes that resumed meiosis and broke down their nuclear envelopes (Fig. 3D). These results indicate that AURKA is essential for meiotic maturation.

To further confirm that AURKB/C cannot compensate for loss of AURKA, we microinjected cRNAs encoding E*gfp* fusions of *Aurka*, *Aurkb*, or E*yfp* fusion of *Aurkc* into *Aurka* KO oocytes. We then visualized bipolar spindle formation and chromosome segregation via live light-sheet microscopy (Fig. 3E-F and Movie S1). As expected, 80% of oocytes from WT mice completed MI, extruded a polar body and reached Met II. In contrast, none of the *Aurka* KO oocytes extruded a polar body and remained arrested at Met I. Exogenously expressed AURKA-EGFP localized to MTOCs and decorated MI spindle poles in *Aurka* KO oocytes. Importantly, AURKA-EGFP expression rescued nearly all *Aurka* KO oocytes because they extruded polar bodies and arrested at Met II (Fig. 3F). Ectopic expression of AURKB-EGFP or AURKC-EGFP, however were unable to rescue MI failure and none of these oocytes extruded a polar body. We were surprised that exogenous expression of AURKC-EGFP could not rescue the ability to complete MI, because a sub-population of AURKC localizes to meiotic spindle poles [20] (Figs. 2E and S1) and the AURKs have some overlapping substrate specificity [34]. This failure to rescue suggests that AURKA and AURKC have unique functions that are likely spatially distinct at the poles.

### *Aurka* KO oocytes are defective in MI spindle building

To determine what unique functions AURKA is required for, we next evaluated spindle formation using immunofluorescence staining of fixed oocytes. Inhibition of AURKA with MLN8237 causes MI spindle defects, ranging from bipolar spindles of reduced length, spindles with multiple poles and to monopolar spindles [8, 26] (Fig. S2A-B). We matured oocytes for the time it took the WT oocytes to reach early pro-Metaphase I (pro-Met I) (3h), late pro-Met I (5h), and Met I (7h) stages *in vitro* prior to fixation (Fig. 4A). We observed differences between oocytes in early pro-Met I. At this first time point, chromosomes in WT oocytes resolved from one another, consistent with the presence of a microtubule ball that makes transient interactions with chromosomes (Fig. 4A-B). The microtubule ball was associated with multiple small, γ-Tubulin-positive MTOCs indicating that MTOC fragmentation occurred [10]. In contrast, the chromosomes in the majority of *Aurka* KO oocytes did not resolve from one another at early pro-Met I (Fig. 4B). There were fewer and larger γ-Tubulin foci indicating a failure to fragment MTOCs. Next, when WT oocytes transitioned from pro-Met I to Met I, the spindles elongated while chromosomes aligned at the Met I plate. Multiple MTOCs fused together to form two well-defined poles. *Aurka* KO oocytes, however, either had a persistent small microtubule ball with unresolved chromosomes or had elongated spindles. Interestingly, both types of spindles always had 1-2 MTOCs that did not fragment and that were not sorted into spindle poles. When we quantified the distribution of these two spindle phenotypes in KO oocytes, ~55% had monopolar spindles, and ~45% had short bipolar spindles after 7h of meiotic maturation (Fig. 4C). We also quantified these spindle phenotypes using length and volume measurements. *Aurka* KO oocytes had significantly shorter bipolar spindles (29.54 μm vs 15.92 μm, WT and KO, respectively) and reduced spindle volume (1219 μm^3^ vs 438.8 μm^3^, WT and KO, respectively) compared to WT oocytes (Fig. 4D-E).

**Fig 4.**
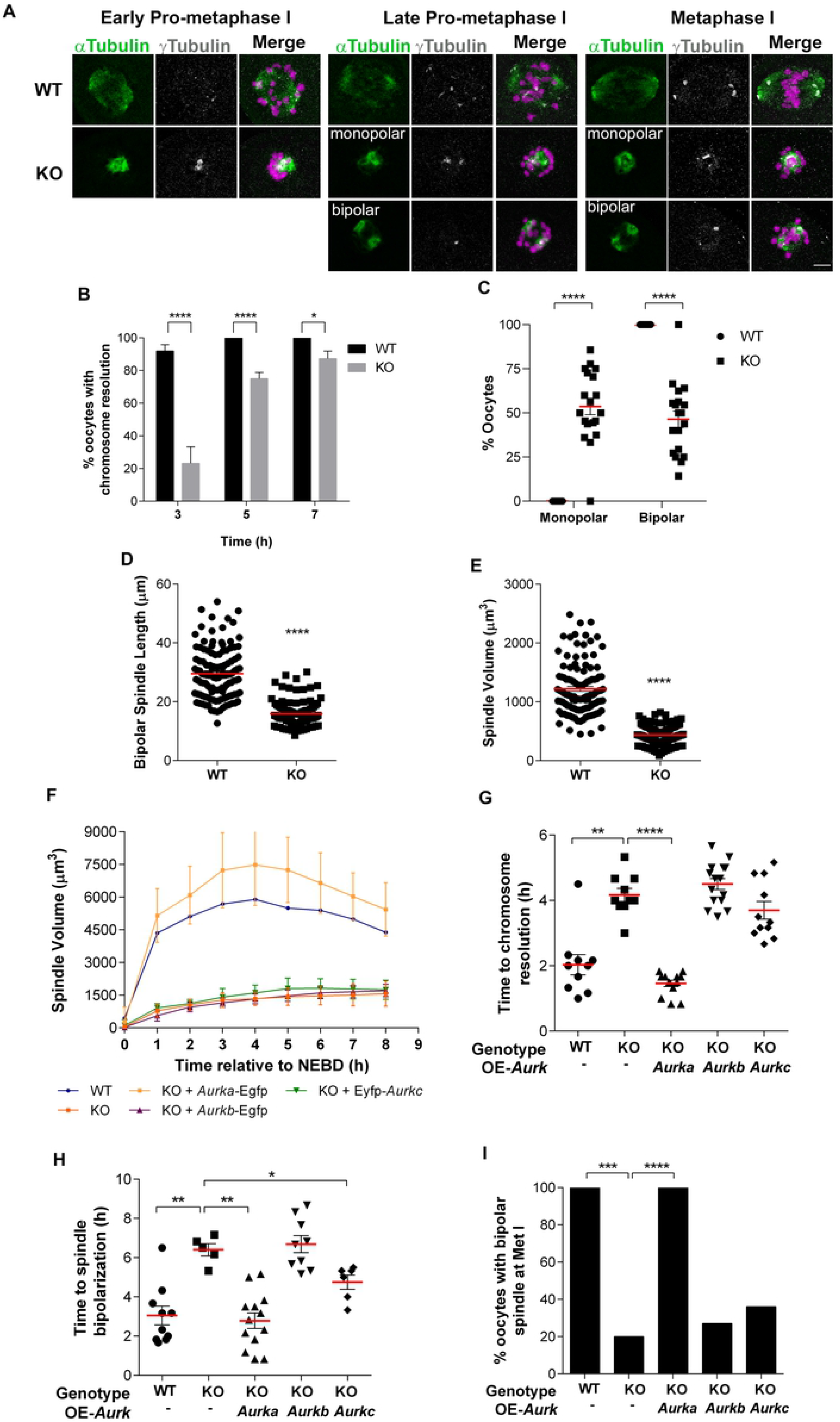
*Aurka* KO oocytes have defects in spindle building. **(A)** Representative confocal images of oocytes from WT and *Aurka* KO females matured to different stages of meiosis, as indicated, and immunostained with antibodies against γ-Tubulin (gray), α-Tubulin (green) and DAPI (magenta). **(B)** Quantification of the percentage (%) of oocytes with resolved chromosomes at different meiotic stages (Unpaired Students t-Test, two-tailed, **** p<0.0001; * 0.0196, 6 experimental replicates). **(C)** Quantification of the % of oocytes with a monopolar or bipolar spindle (Two-way ANOVA; ****p<0.0001, 19 females/genotype). **(D)** Quantification of spindle lengths of bipolar spindles (Unpaired Students t-Test, two-tailed, **** p<0.0001; number of oocytes, WT: 119; KO: 104). **(E)** Quantification of spindle volume (Unpaired Students t-Test, two-tailed, **** p<0.0001; number of oocytes, WT: 113; KO: 143). **(F-G)** image data from Fig. 3E was used for quantification. n (WT, KO, KO + *Aurka*, KO + *Aurkb*, KO + *Aurkc*) = 10, 10, 13, 15, 11, respectively. **(F)** Spindle volume during meiotic maturation. Time is relative to NEBD. **(G)** Time for chromosome individualization and **(H)** spindle bipolarization (Mann-Whitney test, * p<0.05, ** p<0.01, *** p<0.001, **** p<0.0001). **(I)**% of oocytes that had bipolar spindle in Met I (Fisher Exact test, *** p<0.001, **** p<0.0001).

To determine if these spindle defects reflect unique AURKA functions, we used these same spindle quantification parameters to assess if the ectopic expression of each of the Aurora kinases can rescue specific steps of meiotic spindle building. Expression of AURKA rescued all the defects: MI spindle volume was restored, chromosomes resolved from one another with WT-like kinetics, and a stable, bipolar MI spindle formed (Fig. 3E, 4F-J, Movie S1). Expression of AURKB-EGFP failed to rescue all of these parameters. Interestingly, expression of AURKC-EGFP partially rescued the time in which some *Aurka* KO oocytes formed a bipolar spindle (Fig. 4H), although the total number of oocytes that could maintain a bipolar spindle through Met I did not significantly improve (Fig. 4I). Taken together, these results suggest that AURKA is uniquely needed for MTOC fragmentation and building a bipolar MI spindle.

### AURKA is required to fragment MTOCs through PLK1 and to form the LISD

Because our analyses showed that *Aurka* KO oocytes are defective in MTOC fragmentation, we further investigated this phenotype using high-resolution light-sheet microscopy (Movie S2). Oocytes expressed H2B-mCherry, CDK5RAP2-Egfp and were stained with a fluorogenic drug, SiR-tubulin, for visualization of chromosomes, MTOCs, and microtubules, respectively (Fig. 5A). For technical reasons, we started live imaging 40-50 minutes after meiotic maturation began which corresponded to 10-20 minutes prior to nuclear envelope break down in WT oocytes. Typically, at this time in control oocytes, one large dominant MTOC and multiple small MTOCs were present in the cytoplasm and a few smaller MTOCs were in the perinuclear region (Fig. 5A). When oocytes exited prophase I, as marked by nuclear envelope break down and chromosome condensation, the majority of cytoplasmic MTOCs including the dominate MTOC, moved toward the condensing chromosomes. As a result, a microtubule ball formed, and chromosomes resolved. As control oocytes transited from pro-Met I to Met I, MTOCs sorted, spindles elongated and finally MTOCs fused to form two spindle poles. In contrast, and consistent with our previous result (Fig. 4A), *Aurka* KO oocytes lacked multiple cytoplasmic MTOCs and had only 1-2 large MTOCs at the time of meiotic resumption. *Aurka* KO oocytes never fragmented the large MTOC (0/16 MTOC fragmentation in KO vs 12/12 MTOC fragmentation in WT) (Fig. 5A; Movie S2).

**Fig 5.**
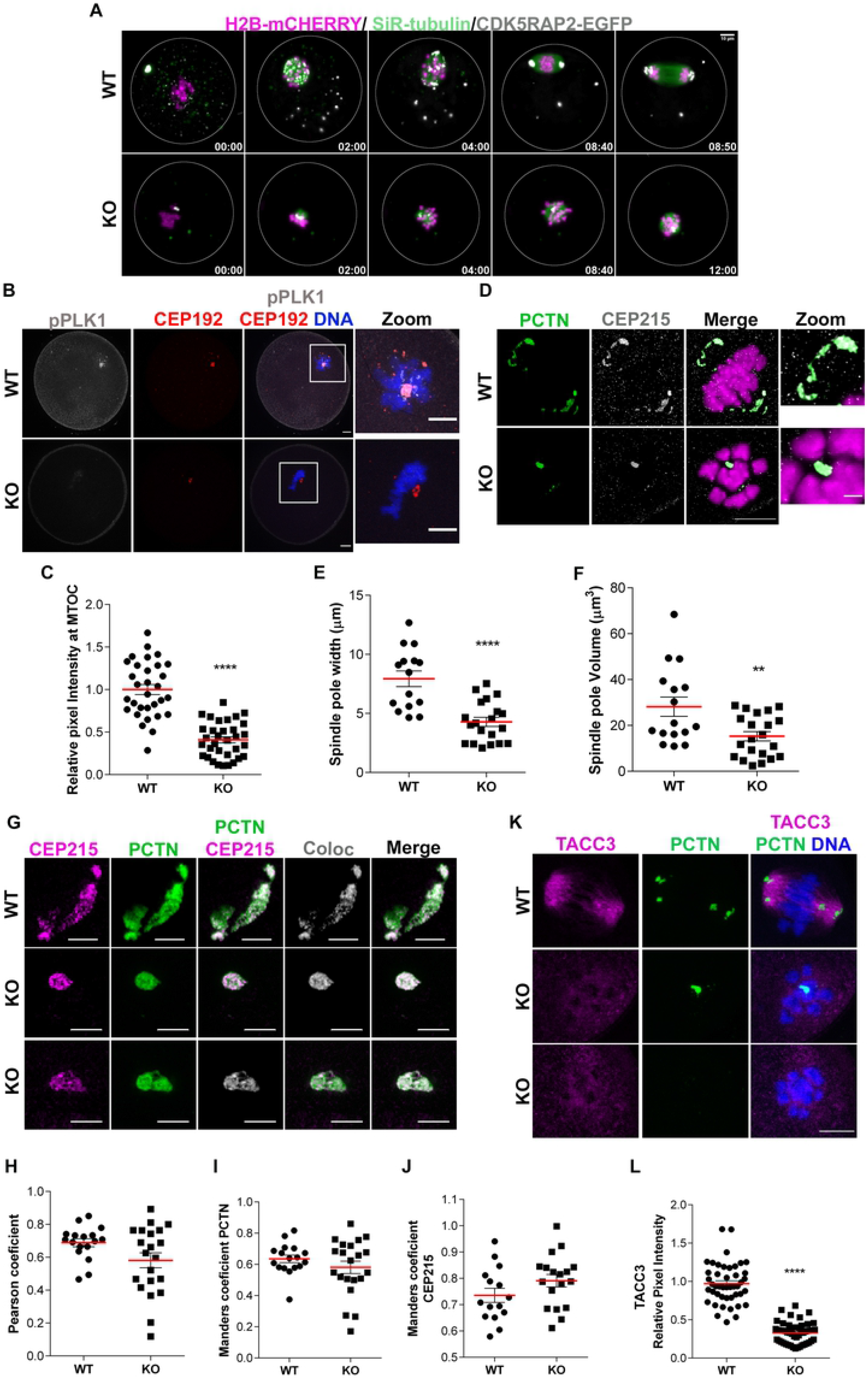
*Aurka* KO oocytes fail to fragment MTOCs and form LISD. **(A)** Representative images of maximum intensity z-projections from WT and *Aurka* KO oocytes matured live using light-sheet microscopy. Oocytes expressed CDK5RAP2-EGFP (MTOCs, gray) and H2B-mCherry (DNA, magenta) while incubated with SiR-tubulin (spindle, green) are shown. Time points are relative to time after nuclear envelope breakdown (h:min). **(B)** Representative confocal images of oocytes from WT and KO females in early pro-Metaphase I immunostained with antibodies against phosphorylated PLK1 (pPLK1, gray), CEP192 (red) and DAPI (blue). **(C)** Quantification of pPLK1 intensity at MTOCs (Unpaired Students t-Test, two-tailed, **** p<0.0001; number of oocytes, WT: 31; KO: 33). (**D)** Representative images of Metaphase I oocytes from WT and KO females visualized with super-resolution microscopy (using Lightning settings) and immunostained with antibodies against Pericentrin (PCTN, green), CEP215 (gray) and DAPI (magenta). Scale bars: 10μm and 2 μm. **(E)** Quantification of spindle pole width in D (Unpaired Students t-Test, two-tailed, **** p<0.0001; number of oocytes, WT: 15; KO: 20). **(F)** Quantification of spindle pole volume in D (Unpaired Students t-Test, two-tailed, ** p=0.0051; number of oocytes, WT: 16; KO: 21). **(G)** Representative images from STED microscopy of spindle poles of WT and KO oocytes at Metaphase I immunostained with antibodies against Pericentrin (PCTN, green), CEP215 (magenta); colocalization specific pixels (gray). **(H)** Quantification of Pearson coefficient. **(I, J)** Quantification of Manders coefficient for Pericentrin (Unpaired Students t-Test, two-tailed, p=0.2736; number of oocytes, WT: 17 A KO: 21) and CEP215 (Unpaired Students t-Test, two-tailed, p=0.129; number of oocytes, WT: 15; KO: 18 respectively. Scale bars: 3 μm. **(K)** Representative confocal images of oocytes from WT and KO females at Metaphase I immunostained with antibodies against TACC3 (magenta), PCTN (green) and DAPI (blue). Scale bars: 10μm and 2μm. **(L)** Quantification of TACC3 intensity (Unpaired Students t-Test, two-tailed, **** p<0.0001; number of oocytes, WT: 45; KO: 49).

Because of the technical limitations of being unable to image live oocytes immediately after induction of meiotic maturation, and because we immediately observed multiple cytoplasmic MTOCs in control but not *Aurka* KO oocytes, we compared the number of MTOCs in prophase I-arrested oocytes after fixation and immunostaining. Both in WT and KO groups we found 1-2 large MTOCs (Fig. S3), suggesting that MTOC defects do not occur during oocyte growth but, instead the first defect in *Aurka* KO oocytes is the inability to fragment MTOCs upon exiting from prophase I.

To understand the role of AURKA in MTOC fragmentation, we evaluated the MTOC regulatory pathway in more detail. Similar to *Aurka* KO oocytes, *Plk1* KO oocytes also arrest in MI with short spindles and have deficiencies in fragmenting MTOCs [35]. Because AURKA can activate PLK1 via phosphorylation of Threonine 210 [36], we reasoned that AURKA functions upstream of PLK1 in mouse oocytes. To test this hypothesis, we performed immunocytochemistry to detect the activated form of PLK1 (pPLK1) in WT and *Aurka* KO oocytes. Consistent with our hypothesis, PLK1-T210 phosphorylation was significantly decreased in *Aurka* KO oocytes at MTOCs (Fig. 5B, C). These data suggest that AURKA regulates MTOC fragmentation by phosphorylating and thereby activating PLK1 after NEBD.

Next, we used super resolution microscopy to understand the consequences of the failure of MTOC fragmentation on Met I spindle pole structure by assessing PCM components pericentrin (PCNT) and CEP215 [26]. WT oocytes had two poles, each of which had the characteristic broad MTOC structure of Met I oocytes. In contrast, *Aurka* KO oocytes had one hyper-condensed spindle pole, with reduced volume and width (Fig. 5D-F). These results are consistent with previous findings that show a collapse of spindle poles after AURKA inhibition [26]. Because we observed changes in the structure of spindle poles in *Aurka* KO oocytes, we used STED-based microscopy to evaluate if AURKA is required for the organization of PCM components. We evaluated the levels of colocalization between CEP215 and PCNT by measuring the covariance in the signal intensity between the two proteins (Pearson coefficient) and by measuring the proportion of overlap of one protein with respect to the other (Manders coefficient). However, we did not observe statistically significant differences in the patterns of colocalization between WT and KO oocytes (Fig. 5G-J), suggesting that the arrangement of these PCM components is not controlled by AURKA.

Recent evidence indicates that the MI spindle has phase-separated structures that aid in its formation [8]. A key component and marker of this liquid-like spindle domain (LISD) is TACC3, a known AURKA substrate [19, 34, 37]. Consistent with this connection, inhibition of AURKA with MLN8237 disrupted the LISD in mouse oocytes. We therefore evaluated if the LISD property was disrupted in *Aurka* KO oocytes. Upon probing WT and *Aurka* KO oocytes with anti-TACC3 antibodies, we found loss of TACC3 signal, corroborating a function of AURKA in coordinating the LISD during MI (Fig. 5K-L). Taken together, these results indicate that AURKA is required to build a proper MI spindle through controlling the initial step of fragmenting MTOCs and formation of a LISD.

### AURKA regulates REC8 cleavage independent of the APC/C

Finally, we evaluated a potential mechanism that would cause the failure to extrude a polar body, despite some oocytes appearing to have small, bipolar spindles. One possibility is the spindle assembly checkpoint (SAC). Insufficient tension between kinetochores and microtubules activates an error-correction pathway involving AURKB/C which triggers detachment of MTs from kinetochores. This loss of kinetochore-microtubule (K-MT) attachments activates the SAC [38] and results in cell-cycle arrest preventing anaphase I. We suspected the arrest in *Aurka* KO oocytes was due to a lack of tension from monopolar and short spindles. We investigated the strength of the SAC in Met I by evaluating MAD2 signals at kinetochores (Fig. 6A). When normalized to kinetochore signal *Aurka* KO oocytes had significantly higher MAD2 than WT oocytes (Fig. 6B). These data suggest persistent SAC activity in KO oocytes, likely due to a defective spindle and the loss of tension.

**Fig 6.**
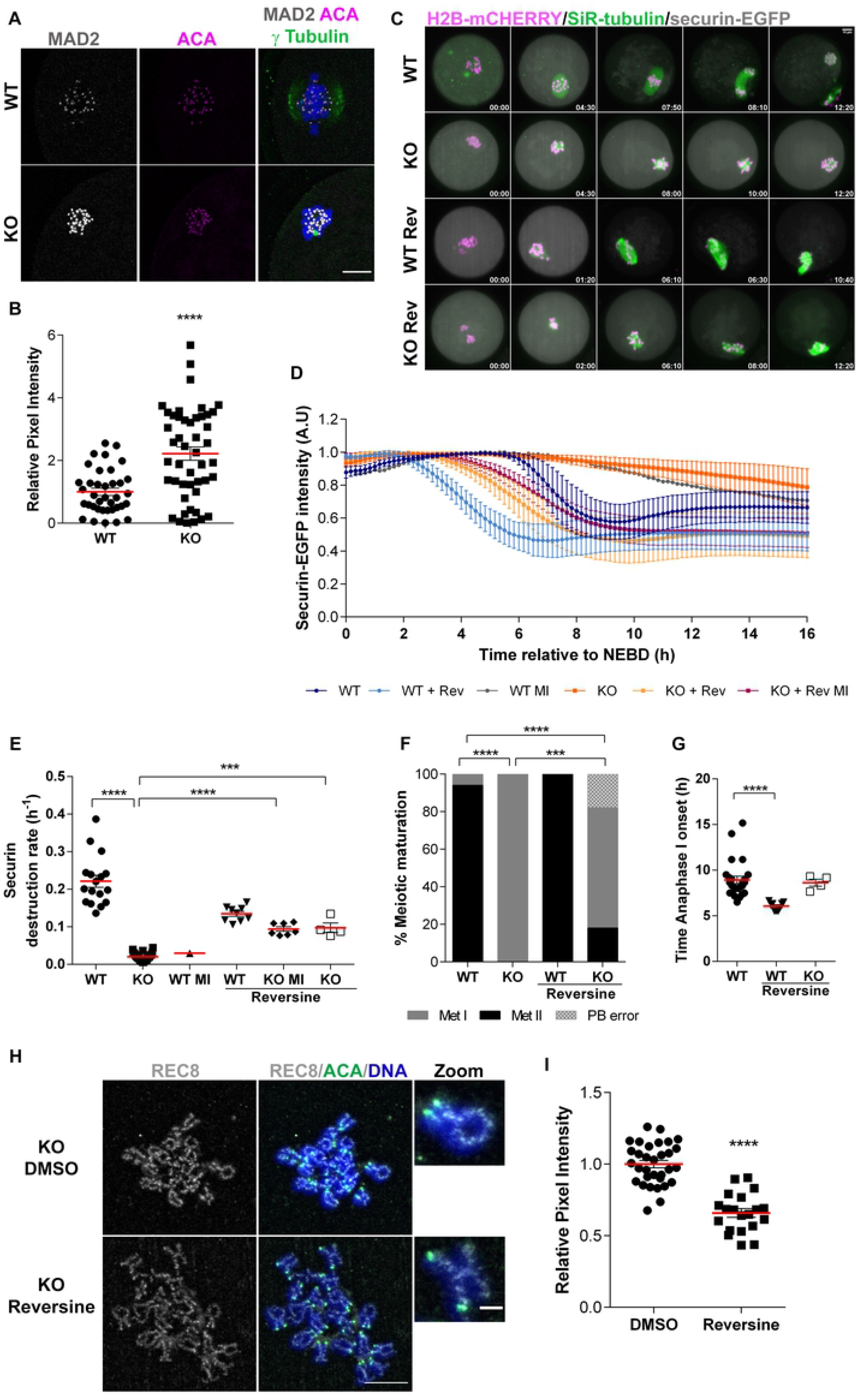
*Aurka* KO oocyte arrest is SAC independent. **(A)** Representative confocal images of oocytes from WT and *Aurka* KO females at Metaphase I immunostained with antibodies to detect centromeres (anti-centromeric antigen (ACA) (magenta)), MAD2 (gray) and chromosomes (DAPI (blue)). **(B)** Quantification of MAD2 intensity at kinetochores in (A) (Unpaired Students t-Test, two-tailed, **** p<0.0001; number of oocytes, WT: 37, A KO: 47). Scale bars: 10μm. **(C)** Live light-sheet imaging of oocytes expressing securin-EGFP (grey), H2B-mCherry (magenta, chromosomes) and stained with SiR-tubulin (green, microtubules) +/− 1μM reversine treatment. Maximum intensity z-projection images are shown. Time relative to NEBD. Scale bar = 10 μm. **(D-G)** Data from (C) was used for analysis. Number of oocytes, WT: 18, KO: 24, WT + reversine: 9, KO + reversine: 11. **(D)** Normalized intensities of cytoplasmic securin-EGFP signals. For normalization, maximum securin-EGFP signal in each oocyte was set to 1. Average +/− SD are shown. **(E)** Rate of securin-EGFP destruction (h^−1^) (Mann Whitney Test, ** p<0.001, **** p<0.0001. **(E)** Proportion of WT and KO oocytes +/− 1μM reversine that reached different phases of meiosis (Met I – metaphase I, Met II – metaphase II, PB error – polar body extrusion retraction) after 16 h of time-lapse imaging (Likelihood Test, *** p<0.001, **** p<0.0001). **(G)** Anaphase I onset (hours relative to NEBD), which was defined as the first time point when segregation of chromosomes was detected (Mann Whitney Test, **** p<0.0001). **(H)** Representative confocal images of chromosome spreads. Oocytes were matured with reversine (1 μM) for 16 h, immunostained with antibodies to detect centromeres (anti-centromeric antigen (ACA) (green), REC8 (gray) and chromosomes (DAPI (blue)). To assess persistent REC8, these oocytes were compared to DMSO-treated KO oocytes. **(I)** Quantification of REC8 intensity in (F) (Unpaired Students t-Test, two-tailed, **** p<0.0001; number of oocytes, KO DMSO: 32 KO reversine: 20). Graphs show the mean ± SEM from at least 3 independent experiments. Scale bars: 10μm and 2μm.

Next, to assess whether the Met I arrest in *Aurka* KO oocytes is solely due to persistent SAC activation, we treated oocytes with reversine to inhibit monopolar spindle 1 (MPS1) kinase, a protein required for initiating the SAC signaling complex [39, 40]. We monitored chromosome segregation, spindle formation and polar body extrusion by light-sheet live imaging (Movies S3-4). As a read-out of Anaphase-Promoting Complex/Cyclosome (APC/C) activity, we also monitored the destruction of securin-EGFP (Fig. 6C). Ninety-five percent of WT oocytes rapidly degraded securin-EGFP (Fig. 6C-E) before anaphase I and extruded the first polar body (Fig. 6F). Anaphase I onset occurred ~9h post-NEBD in this imaging system (Fig. 6G). In contrast, all *Aurka* KO oocytes remained arrested at Met I (Fig. 6F) and had only minor decreases (~10%) in securin-EGFP demonstrating minimal APC/C activity (Fig. 6D-E). Note that in the one WT oocyte that remained arrested in Met I, a similar minor decrease in securin-EGFP also occurred (Fig. 6D-E). As expected, in WT oocytes, reversine-treatment accelerated the onsets of both securin-EGFP destruction (Fig. 6D, E) and anaphase I by 2-3h (Fig. 6G); all oocytes extruded the first polar body (Fig. 6F). To our surprise, although reversine-treatment restored securin-EGFP destruction in *Aurka* KO oocytes (Fig. 6D-E), 64% of these oocytes did not enter Anaphase I and did not extrude a polar body (Fig. 6F). Of the remaining 36% *Aurka* KO oocytes treated with reversine that did enter Anaphase I (Fig. S4A, B; Movie S4), only one-half (18%) extruded the first polar body. The remaining 18% either did not extrude the first polar body or they extruded it, but they then failed cytokinesis and retracted it back into the cytoplasm (Fig. 6F). Importantly, regardless of the polar body extrusion outcome, the APC/C activities in all WT and *Aurka* KO oocytes treated with reversine were similar (Fig. 6E). These data suggest that the Met I arrest in the majority (64%) of *Aurka* KO oocytes treated with reversine cannot be explained by insufficient APC/C activity.

To determine what other functions AURKA may have in controlling anaphase I onset, we speculated that AURKA could also directly regulate chromosome segregation. To undergo anaphase onset, cohesin must be cleaved by separase, which is controlled by APC/C-mediated destruction of securin. When we evaluated the levels of REC8, a meiosis-specific cohesin subunit, in *Aurka* KO oocytes treated with reversine, we found that chromosome-localized REC8 was only reduced by ~ 35% (Fig. 6H-I). These data suggest that AURKA regulates the cleavage of cohesin in an APC/C-independent manner as it does in mitotic cells [41]. Therefore, these data demonstrate that the SAC is not the sole mediator of the Met I arrest in *Aurka* KO oocytes.

In summary, we conclude that AURKA is the only Aurora kinase in mouse oocytes that is essential for fertility and MI [28, 29] (Fig. 7). Its unique functions include initiating MTOC fragmentation through activation of PLK1, spindle formation through regulating TACC3 and the LISD, and anaphase onset through regulating REC8 cleavage. These functions are essential for spindle building and completion of MI to generate a healthy, euploid egg.

**Fig 7.**
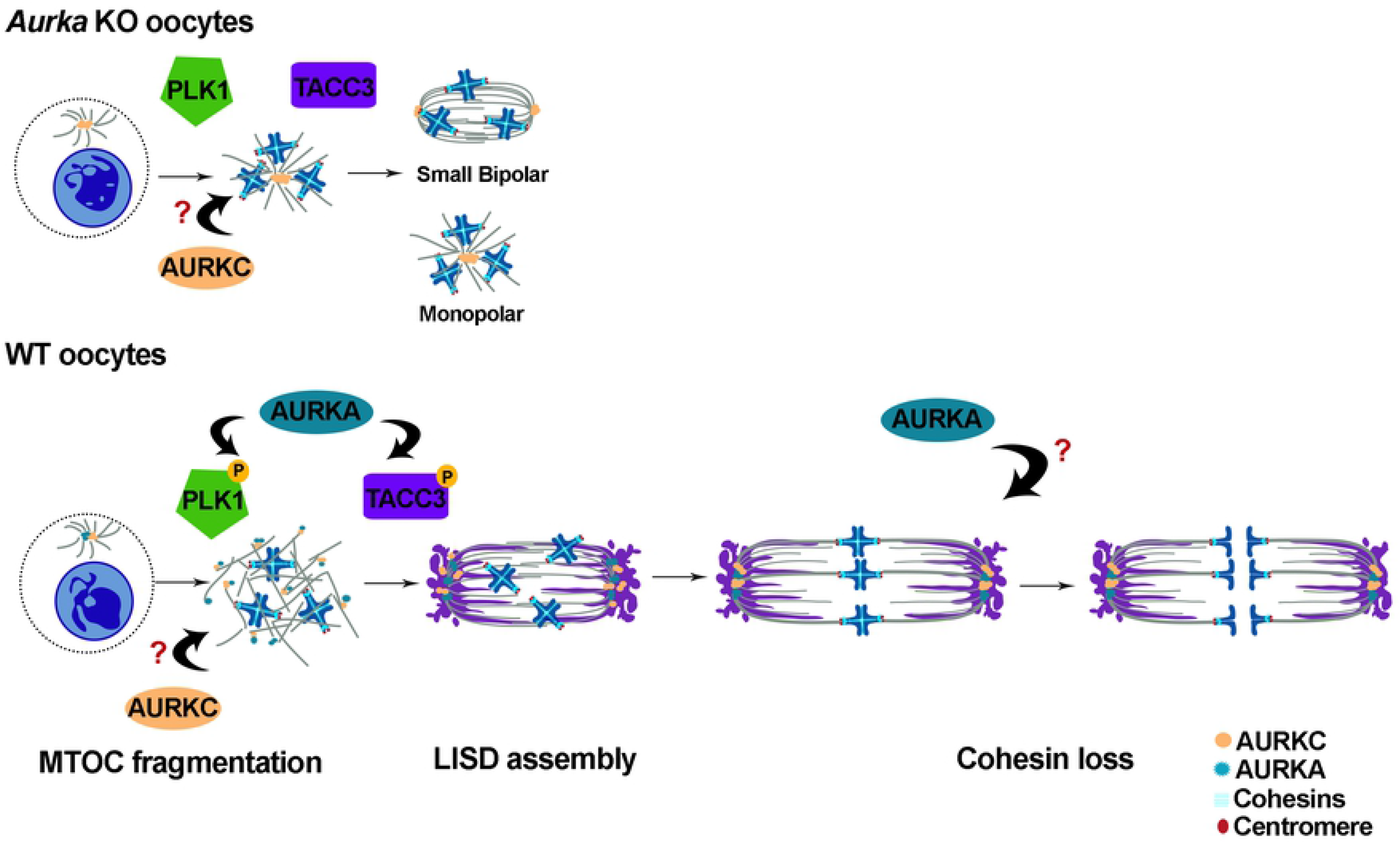
Schematic comparing WT and Aurka KO MI events. In *Aurka* KO oocytes, AURKC still localizes to MTOCs but PLK1 is not phosphorylated and MTOCs fail to fragment. TACC3 does not participate in building the liquid-like spindle domain (LISD). Some spindles are monopolar, but other spindles can become bipolar, but they are short. The result is an MI arrest and a failure to cleave cohesin. In WT oocytes, AURKA and AURKC localize to MTOCs, but likely in distinct regions. AURKA is required to phosphorylate PLK1 to initiate MTOC fragmentation and likely phosphorylates TACC3 to regulate LISD building. Prior to anaphase I onset, AURKA regulates cohesin cleavage in an APC/C independent manner.

## DISCUSSION

Together with our previous description of *Aurkb* and *Aurkc* double knockout oocytes [29], we demonstrate here that AURKA is the only essential Aurora kinase required for mouse female fertility and oocyte meiotic maturation. Although these KO females are sterile, they do ovulate, albeit MI-arrested oocytes. In *Aurkc*^−/−^ *oocytes*, AURKA and AURKB compensate, and in *Aurkb*^−/−^ oocytes, AURKA and AURKC activities are up-regulated. Furthermore, in double *Aurkb*/*Aurkc* knockout oocytes AURKA compensates [28, 29]. Intriguingly, there is no compensatory mechanism for loss of *Aurka*. Specifically, we show that AURKA is needed at the beginning of meiotic resumption for spindle building. AURKA is required for PLK1 activation to initiate MTOC fragmentation and regulates TACC3, likely through phosphorylation, to organize the LISD. Surprisingly, if the SAC is satisfied, AURKA is also required later for Anaphase I onset to trigger efficient cleavage of cohesin by an unknown mechanism (Fig. 7). Collectively, these data imply AURKA-specific substrates or regulatory partner binding that cannot be carried out by the other 2 Aurora kinases.

Substrate phosphorylation by the AURKs is regulated in at least three ways: 1) activation via autophosphorylation, 2) binding to regulator proteins, and 3) phospho-site consensus motifs. Aurora kinase activity depends on T-loop autophosphorylation and binding to regulatory proteins such as TPX2 and INCENP. These regulatory proteins dictate the subcellular localization of the kinases where they can then access their substrates [42–46]. AURKB and AURKC bind INCENP and function in the CPC at chromosomes and kinetochores, whereas AURKA binds MT-binding proteins like TPX2 and functions on spindles and at spindle poles (MTOCs), where AURKA complexes with PCM proteins exist. The binding affinities for these regulatory proteins are governed by the hydrophilicity of an amino acid in kinase subdomain IV [47, 48]. Substitution of this amino acid in AURKA changes the TPX2-dependent polar localization to INCENP-dependent kinetochore localization. This change in localization allows AURKA to compensate in *AURKB*-depleted HeLa cells. Interestingly, the reciprocal residue alteration in AURKB did not facilitate TPX2-binding, and AURKB therefore cannot carry out AURKA function possibly because it cannot activate upon TPX2 association like AURKA can [49]. In our mouse oocyte studies, we observed similar results: AURKA can carry out CPC functions [29], but AURKB/C cannot carry out AURKA spindle pole functions (Fig. 7). Importantly, and different from the HeLa cell experiments, the AURKA-CPC function occurs without amino acid substitution. We speculate that this ability arises because AURKA is the most abundant of the three AURKs in oocytes and there is therefore likely a soluble pool of free AURKA available to bind INCENP when competition is absent.

The third way Aurora kinase phosphorylation is regulation is through sequence specificity for substrates [50, 51]. SILAC-based phosphoproteomics of HeLa cells, revealed that there are many hundreds of AURKA and AURKB substrates and that their phospho-site consensus motifs are similar but distinct. For example, ~91% of AURKA-dependent phospho-peptides contain a R-R-X-p[S/T] motif, whereas only 8% of the AURKB-dependent phospho-peptides contain this motif. Instead, most AURKB-dependent phospho-peptides contained [R/K]-p[S/T]. Therefore, although the motifs are similar, AURKA prefers an arginine at the −2 position and does not prefer a basic amino acid at the −1 position [34]. However, we demonstrated that when AURKA is the sole Aurora kinase in mouse oocytes it can compensate, indicating that AURKA substrate specificity is flexible. This flexibility is consistent with spindle-pole-localized AURKA triggering MT depolymerization at kinetochores [32, 52], a role that AURKB executes at centromeres, and therefore likely through the same protein substrates. In contrast, because AURKB and AURKC cannot compensate for loss of AURKA, even when overexpressed, this consensus-motif flexibility may not be shared. Alternatively, if AURKA occupies part of the LISD, it is possible that the regulatory protein that dictates this particular localization cannot bind and/or activate AURKB/C. Additional subcellular targeting of these kinases could help resolve these mechanistic questions.

Because of the number of possible AURKA substrates, it is likely that KO spindle phenotypes arise from a massive change in substrate phosphorylation and downstream function. For example, we show that PLK1 is not activated in *Aurka* KO oocytes. PLK1 is a known AURKA substrate. PLK1 is required to promote mitotic entry and centrosome maturation through phosphorylation, one substrate being AURKA in a positive feedback loop [53–56]. *Plk1* knockout in mouse oocytes share many phenotypes with the *Aurka* KO oocytes [35]. These phenotypes include sterility, MI arrest with short spindles, and an inability to fragment MTOCs. KIF11 (also known as Eg5) is required for the fragmentation step which occurs after the nuclear envelope breaks down [10], and is a known AURKA substrate in *Xenopus oocytes* [57]. It is therefore likely that a failure to phosphorylate KIF11 can explain the subset of oocytes that retain a monopolar spindle. However, one-half of the oocytes do form bipolar spindles, although they are short. This phenotype suggests that some AURKA-independent MTOC fragmentation can occur, which is not detectable in our imaging system, so that they can form two poles or that chromatin-nucleated microtubules can cluster at their minus ends to form a pole. We also observed short MI spindles in oocytes that lacked *Aurkb/c* where AURKA left the spindle poles and localized to chromosomes to function in the CPC [29]. In mitotic cells, phosphorylation of TPX2 by AURKA is required for MT flux, a function that maintains spindle length [58]. Therefore, in the oocytes with short, bipolar spindles, it is possible that loss of AURKA-TPX2-dependent MT flux has occurred. Moreover, the oocytes with short spindles fail to exit MI even though the APC/C is activated. Another reason of why *Aurka* KO oocytes arrest in MI could be the inability to remove the cohesin proteins that hold homologous chromosomes together. The APC/C releases separase from an inhibitory complex so that it can cleave the cohesin on chromosome arms. Interestingly, the levels of cohesin in *Aurka* KO oocytes where the APC/C was activated were slightly reduced but not completely absent, suggesting that AURKA is involved directly or indirectly in the cleavage of cohesin to trigger Anaphase I onset. In somatic cells, overexpression of AURKA induces loss of cohesin at chromosomes arms by phosphorylating histone H3 at threonine 118 [41]. However, further studies are needed to understand how AURKA regulates cohesin cleavage in mouse oocytes. Finally, another known substrate of AURKA in *Xenopus*, and likely mouse oocytes, is cytoplasmic polyadenylation element binding protein I (CPEB1) [59, 60]. When phosphorylated by AURKA, CPEB1 activates translation of maternal RNAs. In mouse, this burst of translation occurs during oocyte meiotic maturation and is required for completion of MI. Examination of this role in translation in *Aurka* KO oocytes will help explain this cell-cycle arrest.

In summary, we demonstrate that of the 3 Aurora kinases, AURKA is the only essential isoform. This is likely because AURKA regulator partner binding and substrate specificity appear to be more flexible than the other 2 kinases. Because AURKC also localizes to MTOCs, a failure to rescue *Aurka* KO oocytes, even when overexpressed, implies that AURKC is not required for MTOC fragmentation or the LISD, and carries out unknown MTOC and spindle building functions. Identification of MTOC binding partners and substrates will be essential to understanding how AURKA and AURKC coordinate meiotic spindle building.

## MATERIALS AND METHODS

### Generation of mouse strains and genotyping

Mice possessing *loxP* sites flanking exon 2 of the *Aurka* gene [16] were obtained from Jackson Laboratories (B6.129-*Aurka*^*tm1.1Tvd*^/J, #017729). To generate *Aurka*^*fl/fl*^ Gdf9-Cre mice, female mice carrying the *Aurka* floxed alleles were crossed with Gdf9-Cre males (Jackson Laboratories Tg (Gdf9-icre)5092Coo/J, #011062). Animals were maintained following the Rutgers University Institutional Animal Use and Care Committee (Protocol 201702497), National Institutes of Health guidelines, and the policies of the Expert Committee for the Approval of Projects of Experiments on Animals of the Academy of Sciences of the Czech Republic (Protocol 43/2015). These regulatory bodies approved all experimental procedures involving the animals. Mice were housed in 12-12 h light-dark cycle, with constant temperature and with food and water provided *ad libitum*. All animal experiments performed in this study were approved by the Rutgers IACUC.Genotyping for LoxP and Cre were carried out using PCR amplification. Primers for *Aurka LoxP* (Forward: 5’ - CTGGATCACAGGTGTGGAGT- 3’, Reverse: 5’ – GGCTACATGCAGGCAAAC A - 3’), and *Gdf9-Cre* (Forward: 5’ - TCTGATGAAGTCAGGAAGAAC C- 3’, Reverse: 5’ - GAGATGTCCTTCACTCTGATT C-3’, Internal control Forward: 5’ - CTAGGCCACAGAATTGAAAGATCT- 3’, Internal control Reverse: 5’ - GTAGGTGGA AATTCTAGCATCATC C- 3’) were used at 20 pMol using FastMix French PCR beads (Bulldog Bio, #25401) following manufacturer’s protocol.

### Fertility trials

Sexually mature wild-type *Aurka*^*fl/fl*^ and *Aurka*^*fl/fl*^;Gdf9-Cre (*Aurka* KO) female mice ages 5 to 13 weeks were continuously mated to wild type B6D2 (Jackson Laboratories B6D2F1/J, #100006) male mice with proven fertility until a total of 5 litters were produced by WT female mice. Average age of female mice at the end of the fertility trials was 6 months.

### Oocyte collection, culture, and microinjection

Fully grown, prophase I-arrested oocytes were collected from the ovaries of mice ranging in age from 3 to 12 weeks. To prevent spontaneous meiotic resumption during collection, 2.5 μM milrinone (Sigma-Aldrich #M4659) was added to minimal essential medium (MEM). To induce meiotic resumption, oocytes were cultured in milrinone-free Chatot, Ziomek, and Bavister (CZB) [61] medium in an atmosphere of 5% CO_2_ in air at 37 °C. Oocytes were matured for 7.5 hours for metaphase I experiments and 16 hours for Metaphase II experiments.

To obtain oocytes for live light-sheet time lapse imaging, prophase I-arrested oocytes were obtained as described above and oocytes were collected and microinjected in M2 medium (Sigma-Aldrich) and cultured in MEM medium (Sigma-Aldrich) supplemented with 1.14 mM sodium pyruvate (Sigma-Aldrich), 4 mg/ml bovine serum albumin (Sigma-Aldrich), 75 U/ml penicillin (Sigma-Aldrich) and 60 μg/ml streptomycin (Sigma-Aldrich), at 37 °C in a 5% CO2 atmosphere. Oocytes were stained with 100 nM SiR-tubulin (Spirochrome) for microtubule visualization; SiR-tubulin was added to the culture medium. For the securin degradation analysis, a final concentration of 1 μM reversine (Sigma-Aldrich) was added to the oocytes.

For induced ovulation and collection of metaphase II eggs, female mice (>6 wks age) were injected with 5 I.U. of pregnant mare’s serum gonadotropin (PMSG) (Lee Biosolutions #493-10) followed by 5 I.U. of human chorionic gonadotropin (hCG) (Sigma-Aldrich #CG5) 47 h later. 14-16 h post hCG injection, eggs were collected from the oviducts in MEM/polyvinylpyrrolidone media containing 3 mg/ml hyaluronidase (Sigma-Aldrich, #H3506) in MEM for 5 min. Eggs were then washed free of hyaluronidase and allowed to recover in MEM/polyvinylpyrrolidone media prior to fixation.

To inhibit AURKA, MLN8237 (Alisertib, Selleckchem #S1133) was added to CZB culture media at a final concentration of 1 μM. To inhibit the SAC, reversine (Cayman Chemical Research #10004412) was added to CZB culture media at a final concentration of 1 μM. Dimethyl sulfoxide (Sigma Aldrich #472301) was used as a control in the same dilution factor (1:1,000).

After removing the cumulus cells, oocytes were microinjected in M2 medium with ~10 pl of 50 ng/μl *H2b-mCherry*, 125 ng/μl *Egfp-Cdk5rap2*, 100 ng/μl *Aurka-Gfp*, 100 ng/μl *Aurkb-Gfp*, 100 ng/μl *Aurkc-Yfp*, 75 ng/μl *securin-Gfp* cRNAs. Microinjected oocytes were cultured for 3 h in MEM medium supplemented with Milrinone to allow protein expression prior to experimental procedures.

### Plasmids

To generate cRNAs, plasmids were linearized and *in vitro* transcribed using a mMessage mMachine T3 (Ambion #AM1348) and T7 kits (Ambion #AM1344), according to manufacturer’s protocol. The synthesized cRNAs were then purified using an RNAeasy kit (Qiagen #74104) and stored at −80 °C. The pYX-EGFP plasmid was created by transferring T3-T7 cassette from pRNA-EGFP vector [62] into the pXY-Asc vector (NIH, Bethesda, MD, USA) using PCR cloning. The pYX-EYFP plasmid was created from pYX-EYFP plasmid by replacing coding sequence for EGFP by EYFP. AURKC coding sequence [29] was cloned by PCR into pYX-EYFP to create pYX-AURKC-EYFP plasmid. pIVT-AURKB/C-EGFP and pGEMHE-mEGFP-mCDK5RAP2 plasmids were described previously [20, 29].

### Western blotting

A total of 100 prophase-I arrested oocytes were pooled and mixed with Laemmli sample buffer (Bio-Rad, cat #161-0737) and denatured at 95°C for 10 min. Proteins were separated by electrophoresis in 10% SDS polyacrylamide precast gels (Bio-Rad, #456-1036). The separated polypeptides were transferred to nitrocellulose membranes (Bio-Rad, #170-4156) using a Trans-Blot Turbo Transfer System (Bio-Rad) and then blocked with 2% ECL blocking (Amersham, #RPN418) solution in TBS-T (Tris-buffered saline with 0.1% Tween 20) for at least 1h. The membranes were incubated overnight using the antibody dilution anti-AURKA (1:500; Bethyl #A300-072A), or 1 h with anti-MSY2 (1:20,000; gift from R. Schultz) as a loading control. After washing with TBS-T five times, the membranes were incubated with anti-rabbit secondary antibody (1:1000; Kindle Bioscience, #R1006) for 1 h followed with washing with TBS-T five times. The signals were detected using the ECL Select western blotting detection reagents (Kindle Bioscience, #R1002) following the manufacturers protocol. Membranes were stripped prior to loading control detection using Blot Stripping Buffer (ThermoFisher Scientific #46430) for 30 minutes at room temperature.

### Immunocytochemistry

Following meiotic maturation, oocytes were fixed in PBS containing paraformaldehyde (PFA) at room temperature (CREST, α-tubulin: 2% PFA for 20 mins; TACC3, CEP192: 2% PFA for 30 min; phosphorylated PLK1-T210: 2% PFA + 0.1% Triton-X for 20 mins; Pericentrin, phosphorylated CDC25B-S353 and γ-tubulin, CEP215: 3.7% PFA for 1 h), PHEM (PIPES 60mM, HEPES 25mM, EGTA 10mM, and MgCl_2_ 2mM) containing paraformaldehyde (MAD2: 2% PFA for 20 mins) or 100% Methanol for 10 min for AURKA followed by 3 consecutive washes through blocking buffer (PBS + 0.3% (wt/vol) BSA + 0.1% (vol/vol) Tween-20). Prior to immunostaining, oocytes were permeabilized for 20 min in PBS containing 0.1% (vol/vol) Triton X-100 and 0.3% (wt/vol) BSA followed by 10 min in blocking buffer. Immunostaining was performed by incubating cells in primary antibody for 1 h a dark, humidified chamber at room temperature or overnight at 4°C followed by 3 consecutive 10 min incubations in blocking buffer. After washing, secondary antibodies were diluted 1:200 in blocking solution and the sample was incubated for 1 h at room temperature. After washing, the cells were mounted in 5 μL VectaShield (Vector Laboratories, #H-1000) with 4′, 6-Diamidino-2-Phenylindole, Dihydrochloride (DAPI; Life Technologies #D1306; 1:170).

### Chromosome spreads

Chromosome spreads was performed as previously described [63]. *Aurka* KO oocytes were matured *in vitro* for 16 h in DMSO or reversine 1μM and were treated with Acidic Tyrode’s solution (Millipore Sigma; MR-004-D) to remove the zona pellucida. Then, oocytes were transferred to a drop of chromosome spread solution (0.16% Triton-X-100, 3 mM DTT (Sigma-Aldrich, 43815), 0.64% paraformaldehyde in distilled water) on glass slides and allowed to slowly air dry prior to processing for immunofluorescence staining. Immunostaining of chromosome spread was performed by washing the slide two times with PBS for 10 min and blocking the slide in PBS supplemented with BSA 3% for 10 min. Primary antibody to detect REC8 was administered for 3 h in a dark, humidified chamber at room temperature, followed by three washes in PBS of 10 min each. Secondary antibody was applied for 1.5 h in a dark, humidified chamber at room temperature followed by three washes in PBS of 10 min each. After washing, the slide was mounted in Vectashield containing DAPI (Life Technologies, #D1306)

### Antibodies

The following primary antibodies were used for immunofluorescence (IF) experiments: mouse anti α-tubulin Alexa-fluor 488 conjugated (1:100; Life Technologies #322588) AURKA (1:500; Bethyl #A300-072A), ACA (1:30; Antibodies Incorporated #15–234), phosphorylated CDC25B (1:100; Signalway Antibodies #11949), γ-tubulin (1:100; Sigma-Aldrich #T6557), MAD2 (1:100; Biolegend #PRB452C), MSY2 (1:20,000; gift from R. Schultz) [64]. TACC3 (1:100; Novus Biologicals # NBP2-67671), REC8 (1:1000, gift form M. Lampson). Phosphorylated PLK1 (BD Pharmigen #558400); Pericentrin (BD Biosciences, #611814); CEP215 (EMD Millipore #06-1398); CEP192 (Proteintech, #18832-1-AP). The following secondary antibodies were used at 1:200 for IF experiments: Anti-human-Alexa-633 (Life Technologies #A21091), anti-mouse-Alexa-488 (Life Technologies #A11029), anti-rabbit-Alexa-568 (Life Technologies #A10042).

### Microscopy

Images were captured using a Leica SP8 confocal microscope equipped with a 40X, 1.30 N.A. oil immersion objective. For each image, optical z-slices were obtained using a 1.0 μm step with a zoom setting of 4. For comparison of pixel intensities, the laser power was kept constant for each oocyte in an experiment.

To monitor the extrusion of polar bodies, prophase I-arrested oocytes were matured *in vitro* using an EVOS FL Auto Imaging System (Life Technologies) with a 10X objective. The microscope stage was heated to 37°C and 5% CO2 was maintained using the EVOS Onstage Incubator. Images were acquired every 20 min and processed using NIH Image J software.

For super-resolution microscopy we used two different microscopes: a Leica SP8 confocal microscope with Lightning module equipped with a 63X objective, 1.40 NA oil immersion objective. For each image, optical z-slices were obtained using a 0.3 μm step with a zoom setting of 4.5. A Leica SP8 Tau-STED equipped with a 93X objective, 1.3 NA glycerol immersion objective was used to image spindle poles with super resolution. The system was aligned to control any temporal and temperature dependent shift. For each image, optical z-slices were obtained using a 0.17 μm step with a zoom setting of 4.5. Excitation and depletion lasers were kept constant during image acquisition form different genotypes.

Fluorescence time-lapse image acquisitions were performed using Viventis LS1 Live light sheet microscope system (Viventis Miscoscopy Sarl, Switzerland) with a Nikon 25X NA 1.1 detection objective with 1.5 x zoom. Thirty-one 2-μm optical sections were taken with a 750 x 750-pixel image resolution using 10 min time intervals. EGFP, EYFP, mCHERRY and SiR fluorescence were excited by 488, 515, 561 and 638 nm laser lines. EGFP and EYFP emissions were detected using 525/50 (BP) and 539/30 (BP) filters, respectively. For detection of mCHERRY and SiR fluorescence, 488/561/640 (TBP) filter was used.

### Histology

Ovaries of the female mice that were in the fertility trials were fixed in Modified Davidsons fixative solution (Electron Microscopy Sciences, #6413-50) for 6–12 h and were processed by the Office of Translational Science at Rutgers University for histology services. Five μm sections of paraffin embedded ovaries were stained with Harris H/E. Ovarian images were acquired at the 1^st^, 5^th^, and 10^th^ sections in each ovary, under a bright field microscope EVOS FL Auto Imaging System (Life Technologies) with a 20X objective and images were stitched together to project the entire ovary. Ovarian follicles were quantified using morphological criteria [65].

### Image analysis of fixed oocytes

Image J software was used to process most of the images (NIH, Bethesda, USA). For analysis, z-slices for each image were merged into a projection. Bipolar spindle length was measured between the two furthest points on both spindles using the line tool in Image J. Spindle volume was determined using the 3D reconstruction tool in Imaris software (BitPlane) freehand tool to mark precisely around the spindle. For pixel intensity analyses the average pixel intensity was recorded using the measurement tool. To define the region of the chromosomes for intensity measurements, the DNA channel (DAPI) was used as a mask. MTOC markers, including AURKA and γ-tubulin were used to define spindle poles, and CREST was used as a kinetochore marker for pixel intensity measurements. Imaris software we used for colocalization analysis of CEP215 and Pericentrin (BitPlane). Briefly, we determined a region of interest around the spindle pole, we set threshold for each channel and using the colocalization module we determined: Pearson coefficient which measures the covariance in the signal levels of two images; and Manders coefficients which are indicators of the proportion of the signal of one channel with the signal in the other channel over its total intensity [66, 67].

### Image analysis of live oocytes imaged by light-sheet microscopy

All image analysis was done using Fiji software [68]. For analysis of securin-EGFP degradation of the mean intensity of securin-EGFP was measured on a non-signal adjusted middle optical stack in every time frame. In every oocyte, measured mean values from each time point were normalized to the time frame with a maximum mean intensity. Calculation of destruction rate was described previously [69]. Briefly, destruction rate of securin-EGFP (h-1) was defined as the negative value of the slope of the line that can be fitted to the decreasing region of securin-EGFP destruction curve.

### Statistical analysis

T-test and one-way analysis of variance (Anova) were used to evaluate the significant difference among data sets using Prism software (GraphPad Software). The details for each experiment can be found in the Results section as well as the figure legends. “Experimental n” refers to the number of animals used to repeat each experiment. Data is shown as the mean ± the standard error of the mean (SEM). P < 0.05 was considered significant. All statistical analysis of data from live light-sheet microscopy was done using NCSS 11 software (NCSS, LLC; Utah, USA). The type of test used are indicated in the figure legend.

## Acknowledgements

The authors thank Dr. Philip Jordan for assisting with acquisition of the conditional KO mice, Ms. Marianne Polunnas for processing the ovarian histology and Dr. Jessica Shivas for STED imaging acquisition. They acknowledge Drs. Michael Lampson and Richard Schultz for the REC8 and MSY2 antibodies and members of the Schindler and Solc labs for helpful discussions. This work was supported by an NIH grant to KS (R01 GM112801), the Inter-Excellence Program award (LTAUSA17097) to PS, and by the award from National Sustainability Program of the Czech Ministry of Education, Youth and Sports (LO1609).

## Contribution

KS and PS conceived of the project, analyzed data, wrote and edited the manuscript; CB, PI, MV and DD conducted experiments and analyzed data; CB and PI wrote and edited the manuscript; DD and MV edited the manuscript.

## Supplementary figure legends

**Figure S1. AURKC localizes to MTOCs**

Live light-sheet imaging of KO oocytes expressing histone H2B-mCHERRY (magenta), AURKC-EYFP (gray) and stained with SiR-tubulin (green). The arrows point to AURKC localization. Maximum intensity z-projections at Metaphase I. Scale bars: 10 μm.

**Figure S2. Inhibition of AURKA causes spindle defects**

**(A)** Representative confocal images of oocytes at Metaphase I matured with MLN8237 (MLN) and immunostained with antibodies against α-Tubulin (green) and DAPI (gray). **(B)** Quantification of the percentage (%) of oocytes with different spindle phenotypes (Unpaired Students t-Test, two-tailed, * p=0.014). **(C)** Quantification of the bipolar spindle area (Unpaired Students t-Test, two-tailed, **** p<0.0001; number of oocytes, WT: 31; KO: 23). **(D)** Quantification of the bipolar spindle length (Unpaired Students t-Test, two-tailed, **** p<0.0001; number of oocytes, WT: 30; KO: 22). Graphs show the mean ± SEM from at least 3 independent experiments.

**Figure S3. *Aurka* KO oocytes have normal number of MTOCs at prophase I.**

Representative confocal images of WT and *Aurka* KO prophase I-arrested oocytes immunostained with γ-Tubulin (magenta), α-Tubulin (green), DAPI (blue). Scale bar: 20μm.

**Figure S4. Comparison of securin destruction in *Aurka* KO oocytes treated with reversine. (A)** Live light-sheet imaging of KO oocytes expressing securin-EGFP (grey), H2B-mCherry (magenta, chromosomes) and stained with SiR-tubulin (green, microtubules) treated with 1μM reversine. Maximum intensity z-projection images of KO oocyte arrested at MI (KO MI), KO oocyte entering Anaphase I and extruding of polar body (KO MII), and KO oocyte entering Anaphase I but having a polar body emission error (KO PB error). Time relative to NEBD. Scale bar = 10 μm. **(B)** Normalized intensities of cytoplasmic securin-EGFP signals. WT, KO and KO + Reversine MI groups are same as in Fig. 6D. KO + Reversine and KO + Reversine PB error are split from KO + Reversine group in Fig. 6D.

## Supplemental Movie legends

**Movie S1:** Movie corresponding to oocytes presented Fig. 3E.

**Movie S2:** Movie corresponding to oocytes presented Fig. 5A.

**Movies S3:** Movie corresponding to oocytes presented Fig. 6C.

**Movies S4:** Movie corresponding to oocytes presented Fig. S4A

